# The dynamics of hinge flexibility in receptor bound immunoglobulin E revealed by electron microscopy

**DOI:** 10.1101/2023.02.17.528943

**Authors:** Rasmus K Jensen, Michaela Miehe, Rosaria Gandini, Martin H Jørgensen, Edzard Spillner, Gregers R Andersen

## Abstract

Immunoglobulin E is a mammal specific antibody isotype supporting the immune response against parasites and venoms, but also a driver of allergic responses. Prior studies have defined the conformation of the IgE Fc fragment bound to the cell surface receptor FcεRIα and the dynamic properties of the IgE Fc. It remains unknown, how these prior studies translate to the complex of a full antibody including the Fab arms with the receptor. Here we show that in a cryo-EM structure of the IgE FcεRIα complex, IgE adopts a T-like conformation where the antigen binding Fab arms may be parallel to the cell membrane. Two additional conformations are captured in negative stain EM (ns-EM) where the arrangements of the Fab arms differ from the cryo-EM conformation. Small angle scattering data favors the FcεRIα bound IgE conformation observed by cryo-EM, but the major IgE conformation observed by ns-EM possibly may also occur. In all observed conformations of FcεRIα bound IgE, one Fab arm is fixed relative to the IgE Fc moiety whereas the second Fab may alternate its position. Introduction of flexibility in the Fab-Fc hinge diminishes the biological activity of IgE demonstrating a functional role for the observed defined Fab-Fc hinge conformations. Our data show the organization of a full size antibody on its receptor and reveal a new layer of dynamics in FcεRIα bound IgE on top of the well established spectrum of IgE Fc conformations. Development of novel anti-IgE therapeutics may take into account these distinct FcεRIα bound IgE conformations.

**Significance statement:** IgE represents a canonical antibody isotype and is a key molecule for the allergic immune response to environmental triggers driven by mast cells and basophils. The requirements for efficient mediation of IgE’s effects are not fully understood. Here we elucidate the structure of the entire IgE in complex with its high affinity receptor and identify two clearly distinct and dominant conformations, in which one of the Fab arms is fixed relative to the Fc domains. Enforcing IgE flexibility impacts the biological function with potential consequences for the allergic response. This unique behavior makes IgE different from all other isotypes and its understanding sheds light on the allergenic activation of the immune response.

## Introduction

Immunoglobulin E (IgE) has a key role in allergic diseases and is found both in the circulation and on a variety of cell types (1). It interacts directly with the high-affinity receptor FcεRI and the low-affinity receptor CD23 (2). Upon crosslinking by allergens, FcεRI-bound IgE on mast cells and basophils triggers degranulation, release of mediators, and immediate reactions (2) (Fig. 1A).

**Figure 1.**
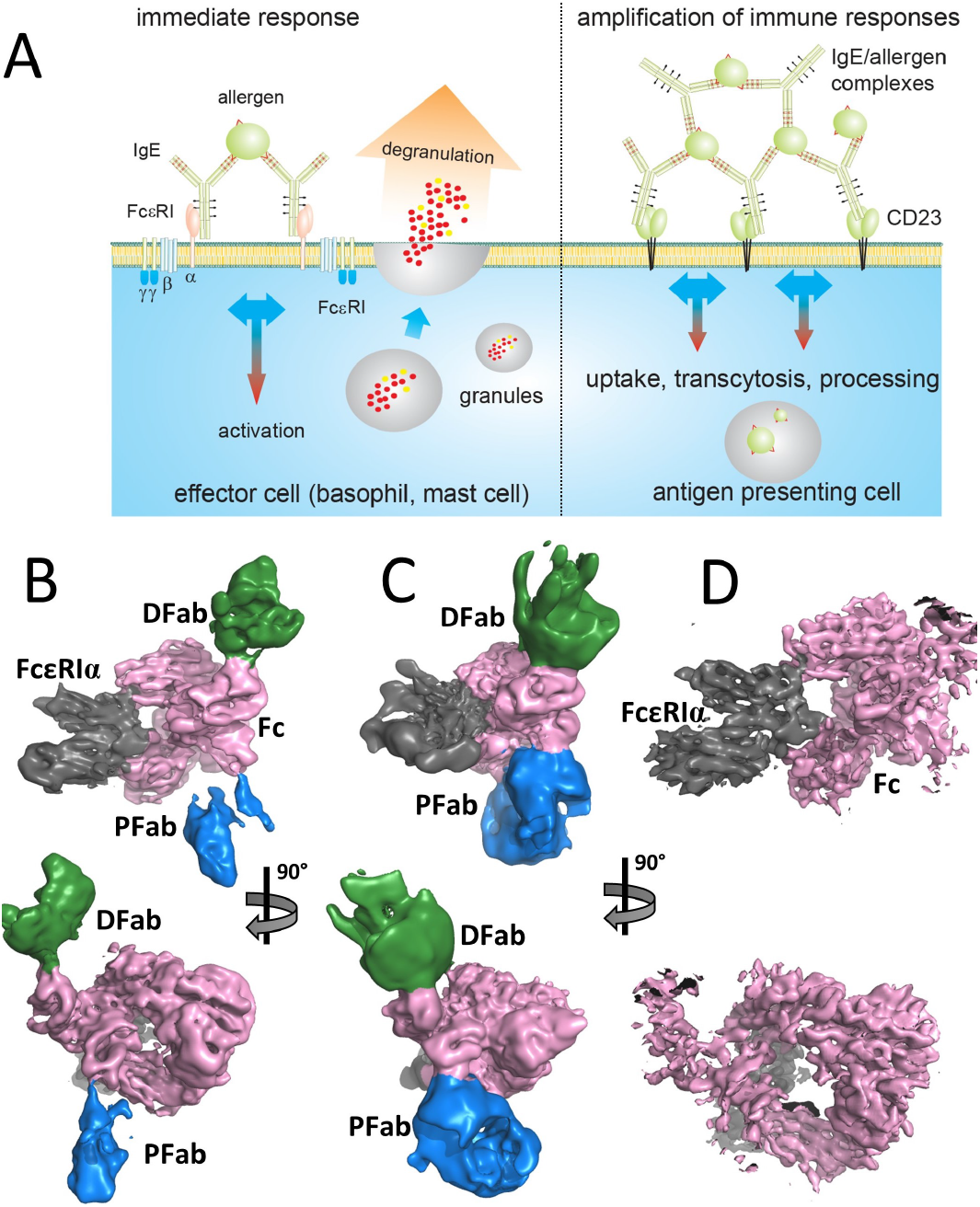
Cryo-EM structure determination of the IgE-FcεRIα complex. A) Representation of the interplay and functional effects of IgE bound to its receptors FcεRI on basophils and mast cells (left) and CD23 on antigen presenting cells (right). B) Two orientations of the consensus cryo EM map of the IgE-FcεRIα complex contoured at 16 σ. Density around the receptor and the IgE Fc is grey and pink, respectively, whereas density at around the distal Fab (dFab) is green and density around the proximal Fab (pFab) is blue. C) As in panel B, but displaying a map filtered by local resolution and contoured at 6 σ to provide a better impression of the Fab arms. At the proximal Fab, the cleft between the variable the constant domains is visible. D) Map from the focused refinement centered on IgE-Fc and FcεRIα contoured at 20 σ.

Crosslinking of IgE-FcεRI complexes by antigens (which may be allergens) induces FcεRI signal transduction through receptor clustering (3). The cytoplasmic domains of the FcεRI β- and γ-subunits contain ITAM motifs that are phosphorylated upon FcεRI crosslinking and link to downstream signaling pathways (4). The homotrimeric CD23 is expressed by epithelial cells, B cells and myeloid cells (5) and is crucial for facilitated antigen presentation, transport across the epithelium, amplification of IgE production and epitope spreading (2, 6) (Fig. 1A).

Antibodies are commonly perceived as flexible molecules due to the presence of a hinge region connecting the antigen recognizing Fab fragment to the constant Fc fragment (7). But in IgE, the hinge-region is short compared to IgG and furthermore the IgE epsilon heavy chain contains four constant domains as compared to three constant domains found in IgG. The short hinge region in IgE reduces flexibility compared to IgG (8, 9) and experimental evidence for limited rotational freedom at the Fab-Fc hinge in both free and FcεRI bound IgE was demonstrated early on by spectroscopy (10) and the formation of 2D crystals of IgE on hapten covered mono- and bilayers (11). There is, however, an understanding of the IgE Fabs as being somewhat flexible relative to the Fc (12-14). A number of structural studies have investigated the conformational properties of the IgE Fc, but little structural work has been done on the full IgE molecule. We recently showed that at the resolution of negative stain EM (ns-EM), the Fabs appears to be rigidly associated with the Fc in intact free IgE and in IgE bound to the monoclonal antibody ligelizumab (15). Prior structural studies using the Fc fragment as a proxy for the full IgE have provided crucial information regarding the fundamental intermolecular interaction between antibody and receptor but these structures do not account for the Fab arms and their impact on the FcεRI bound full antibody (16, 17).

## Results

### Cryo-EM analysis reveals a T-shaped IgE in complex with FcεRIα

To determine an atomic structure of the full IgE bound to the FcεRI receptor, we prepared the complex between the 170 kDa IgE HMM5 and the 50 kDa FcεRIα ectodomain. Cryo-EM grids prepared using standard methods led to strong preferential orientation of the complex, presumably due to interaction with the air-water interface (18). To overcome this complication, we prepared samples on self-wicking Quantifoil Active grids using the Chameleon device (19). Though preferential orientations were still present, we were able to resolve the structure to an overall resolution of 4.1 Å (Fig. 1B-C and Fig. S1A-C). In this consensus map extending over the full complex, the two Fab arms are positioned asymmetrically with respect to the Fc and the receptor (Fig. 1C), with a proximal Fab located close to the Fc, whereas the distal Fab is more protruding.

In contrast to the Fab arms, the IgE Fc and its interface with FcεRI is well resolved and focused refinement on these domains yielded a 3D reconstruction with a resolution of 3.8 Å (Fig. 1D and Fig. S1D). An atomic model for the IgE-FcεRI complex was first obtained by model building and refinement against the focused map which allowed side-chain level modelling in most regions (Fig. S2A-D). The model was subsequently extended using the consensus map to contain the constant domains Cε1 in the heavy chain and CL in the light chain of the two Fab arms. For both Fab-Fc hinges, there is density for the linker between the Cε1 and Cε2 domains (Fig. S3A). The densities for the two Cε1 domains are less well-resolved than their connected Cε2 domains and the local resolution of the Cε1 and the associated CL domains is 7-10 Å (Fig. S1E).

This limited resolution prevented de novo chain tracing of the Cε1 domain for which there is no experimental structure. We therefore generated an alphafold2 model of the HMM5 Fab that as expected matches well with our crystal structure of this Fab where the Cε1 domain was replaced by a Cγ1 domain (20). This Fab model features the disulfide bridge pattern annotated in the uniprot entry IGHE_HUMAN (P01854) and is compatible with the presence of three Asn-linked glycans in the Cε1 domain (Fig. S4A). Due to the limited resolution, the location of the structurally homologous Cε1 and the CL domains could not be unambiguously derived directly from the 3D reconstruction, but comparison of the two possible orientations of the Fab model with the map strongly favored one of these orientations (Fig. 2A and Fig. S3B). In agreement with the selected orientation of the Cε1-CL domain tandem, there is excess density around especially Asn137 and Asn165 compatible with the above mentioned Asn-linked glycans in the Cε1 domain (Fig. S3C). This orientation was therefore selected for model building and refinement against the consensus map.

**Figure 2.**
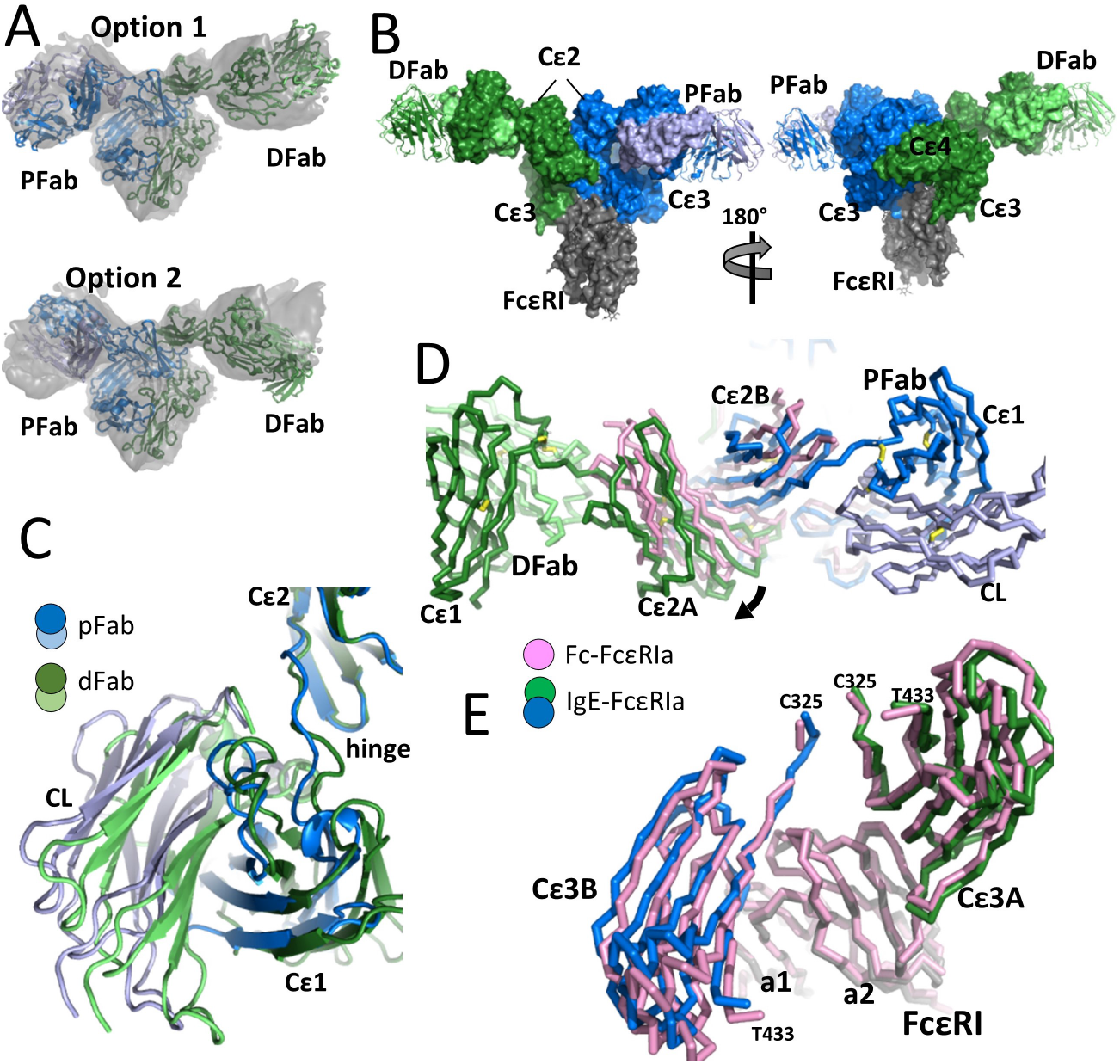
Cryo-EM reveals an IgE T conformation in complex with FcεRIα. A) Superposition of the HMM5 crystal structure in the two possible orientations favors option 1 for distinguishing Cε1 and CL. B) The final structure of the IgE-FcεRIα complex. The Fab VL and VH domains are shown as cartoon as they are not present in the deposited structure. C) Comparison of the two hinge regions and their connected Cε1-CL domain tandems. The two ε-chains are superimposed on their Cε2 domains. D) A superposition of the Fc-FcεRIα and IgE-FcεRIα structures on the receptor reveals a difference in the location of the Cε2 tandem. E) Close-up after superposition on the receptor demonstrating a minor difference in rotation of the Cε3 domains between the Fc-FcεRIα and IgE-FcεRIα structures.

The deposited model fitting the consensus map (Table S1) contains the entire complex except for the variable domains of the Fab arms. The Fab elbow angle measures the angle between the two pseudo-dyad axes relating the two variable domains and the two constant domains, respectively (21). In the crystal structure of the HMM5 where the Cε1 domain is replaced by a Cγ1 domain, the elbow angle is 136°. By measuring the correlation between the consensus 3D map and molecules in a library of 2468 known Fab structures, we found that the Fab structure in the protein data bank (pdb) entry 4dw2 with an elbow angle of 120° fitted the 3D map for both Fab arms better than any other structure in the library. Consequently, an optimized model of the full IgE bound to FcεRIα was created by superposition of the variable domains of the HMM5 structure onto the variable domains in the 4dw2 structures fitted at each Fab arm (Fig. 2B and Fig. S3B). Both Fab arms are stably associated with their respective Cε2 domain in two distinct orientations that differ by 12° (Fig. 2C). The overall appearance of the IgE in the complex resembles that of a slightly distorted T, and if the C-terminal end of the FcεRIα ectodomain is perpendicular to the cell surface, the Fab arms and the Cε4 domain tandem will roughly be parallel to the membrane (Fig. 2B).

Since the distal Fab appears not to interact stably with Cε3-4 domains, it must primarily be the hinge residues and the surrounding residues in the Cε1, CL and Cε2 domains that controls the spatial relationship between the Fab arms and the Cε2 domain tandem. At the proximal Fab, the linker connecting the VH and the Cε1 domains is close to His381 in the Cε3 domain of the same ε-chain. At a low contour level in the consensus map, a density connecting these regions is visible (Fig. S3B). Such additional weak direct contacts formed between the proximal Fab and the Cε3 domain offer an explanation for the difference between the two Fab-hinge-Cε2 conformations. Without the putative Cε3 contact, the hinge conformation at the distal Fab is probably favored energetically over the hinge conformation at the proximal Fab.

### The IgE Fc conformation in complex with FcεRI

Prior structures of the FcεRIα ectodomain in complex with the Cε3-4 and the Fc fragments of IgE revealed that the receptor binding site has contributions from both Cε3 domains and that IgE Fc adopts an asymmetric bent conformation with the Cε2 domain tandem placed oppositely of the FcεRI (16, 17). Comparison of our cryo-EM structure with the crystal structure of the IgE Fc-FcεRI complex demonstrates that the overall features of the receptor-Fc complex with the two interaction sites Cε3A and Cε3B are preserved in the receptor-IgE complex (Fig. S4B). The ε chain recognized by the FcεRIα Cε3A site contains the Cε1 domain of the distal Fab, whereas ε chain in the Cε3B site contains the Cε1 domain of the proximal Fab.

An important descriptor for the overall conformation of the IgE Fc is the bending of the Cε2 domain tandem relative to the two-fold symmetry axis relating the two Cε3 and Cε4 domains. In the IgE containing cryo-EM structure, the Fc bending is 107° compared to 110° in the IgE Fc complex with the FcεRIα ectodomain (17). When the two structures are superimposed on FcεRIα, the orientation of the Cε2 tandem differ by a coordinated 6° rotation relative to the FcεRIα (Fig. 2D). Based on the crystal structures of IgE Fc and its complex with FcεRIα, it was earlier reported that a β-strand shift occurred within the Cε2 domain between the two β-sheets upon complex formation with the receptor (17). Our maps do not support such a strand shift (Fig. S2E-F), and the topology of Cε2 domain in our model of FcεRIα bound IgE is that observed in crystal structures of unbound IgE Fc (22).

In addition to the bending of the Cε2 domain tandem, the Fc Cε3-4 sub-fragment can adopt multiple conformations ranging from closed to open depending on the spacing of their Cε3 domains and Cε3 distance to the Cε4 domains (23). This conformational flexibility allows the Cε3 domains to rotate relative to each other. FcεRI and CD23 recognize different conformations of the IgE Cε3-4 domains, the open and closed state, respectively. In our cryo-EM structure, the Cε3-4 domains are in the open conformation as in the Fc-FcεRI crystal structure but the relative orientation of the two Cε3 domains differ by 6° rotations compared to the Fc containing crystal structure (Fig. 2E). In summary, there are small but significant conformational changes within the Fc part in the cryo-EM structure of full IgE in complex with the FcεRIα ectodomain compared to the crystal structure of the complex containing only the Fc fragment. These differences are likely to be caused by the presence of the Fab arms in the cryo-EM structure, but lattice interactions may also have influenced the conformation of IgE Fc in the crystal structure (17).

### Negative stain EM captures alternative conformations of the receptor bound IgE

Since ns-EM was used for sample quality control, we also obtained two ns-EM 3D reconstructions of the HMM5 IgE-FcεRIα complex (Fig. 3A and Fig. S5). The major 3D class was based on twice as many particles as the minor 3D class. A model of the IgE Fc and the bound receptor ectodomain with the proximal Fab omitted could be fitted convincingly into one of the two hands of each 3D class whereas the model fitted less well in the opposite hand of the two 3D classes (Fig S5D). To complete the models of IgE-FcεRIα complex fitting the ns-EM 3D classes, the proximal Fab was fitted as a rigid body into the density next to the proximal Cε2 domain. Hence, the resulting complete models are obtained by fitting of only two rigid bodies derived from the cryo-EM structure and fit their respective ns-EM 3D reconstructions with correlation coefficients of 0.90 (major) and 0.87 (minor).

**Figure 3.**
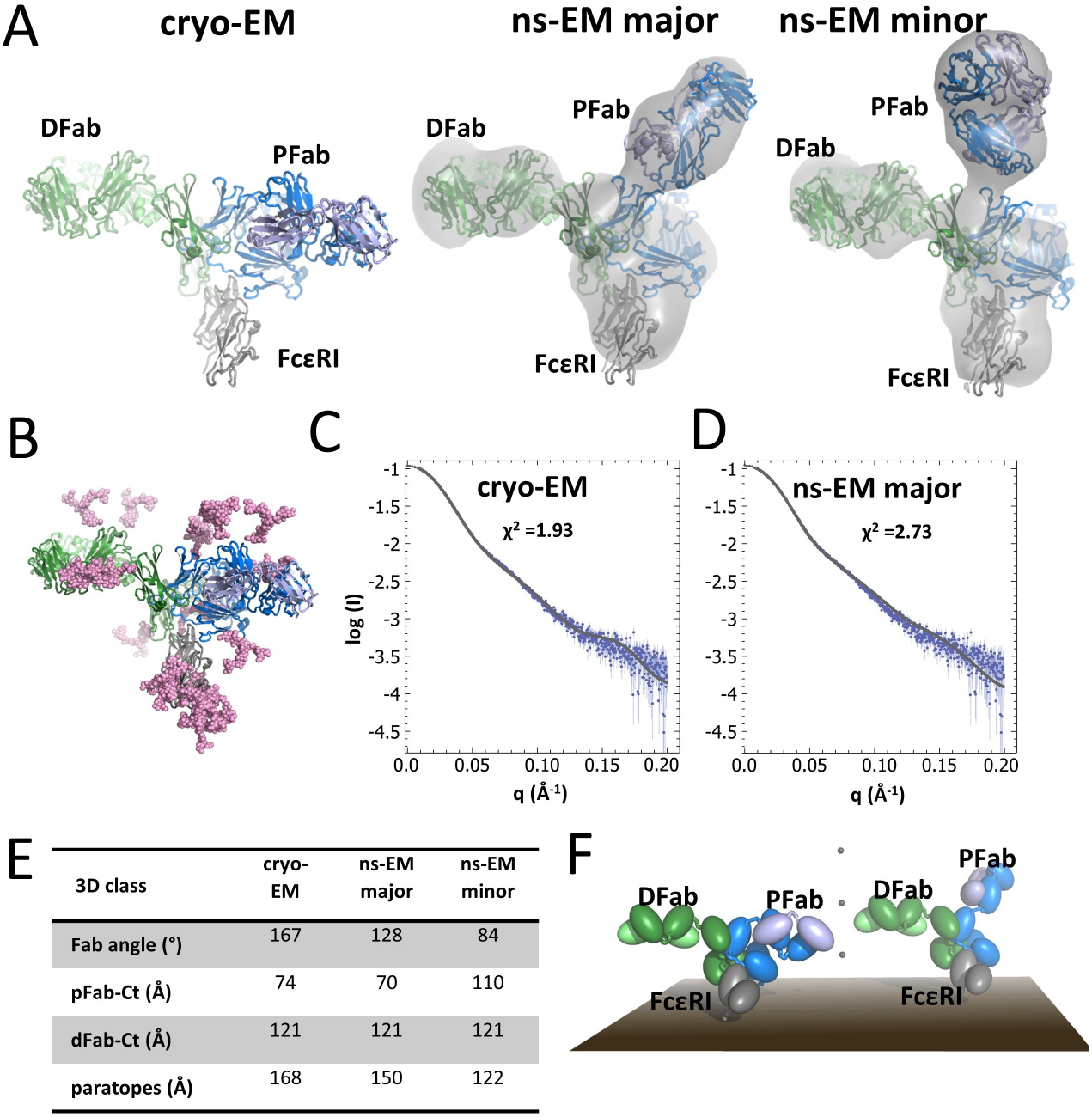
FcεRI bound IgE also adopts non-T conformations. A) At the top, the fitted models of the IgE-FcεRIα complex are shown docked into the two different low resolution ns-EM 3D reconstructions (grey envelope). Notice that the Fab heavy and light chains cannot be distinguished in the ns-EM maps. The cryo-EM based model of the IgE-FcεRIα complex is presented for comparison. B) Model of the fully glycosylated complex in the cryo-EM conformation obtained by SAXS rigid body modelling. The orientation is that of panel A. Glycans are shown as pink spheres. C) Fit of the model in panel B to the experimental data. D) As panel C, but for the fully glycosylated model in the major ns-EM conformation. E) Summary of key geometric properties of IgE in the three conformations presented in panel A. The Cα atoms of heavy chain Ser100 and Ala542 were used to determine the distances (paratope to the C-terminal end of the heavy chain) and (paratope-paratope). F) Models of cell bound IgE-FcεRI complexes illustrates that the one of the IgE paratopes in the ns-EM major class conformation (right) is likely to extend significantly further out from the cell surface compared to IgE in the conformation observed by cryo-EM (left). The three vertically aligned dots in the center are placed 50, 100 and 150 Å above the surface.

In the two ns-EM 3D classes, there are well defined densities for both Fab arms and the Fab arms are also clearly visible in the 2D classes (Fig. 3A and S5A). Hence, the ns-EM 3D reconstructions give further support to the concept that in FcεRIα bound IgE, the Fab arms are fixed relative to the Fc like in unbound IgE (15). However, the orientations of the Fab arms relative to the IgE Fc in the ns-EM 3D maps are clearly different from those present in the cryo-EM structure (Fig 3A).

These differences raised the concern that receptor bound IgE could change conformation during preparation of the EM grids. We therefore conducted a synchrotron SEC-SAXS experiment in which the scattering from the IgE-FcεRIα complex eluting from a size exclusion column was continuously recorded. Data analysis suggested a complex in solution with a maximum extent of 17 nm (Fig. S6A-C). To enable a direct comparison between the observed EM conformations, we calculated scattering curves from molecular models fitted into the two 3D reconstructions. These showed significant deviations from the observed scattering curve and χ^2^ values of 14-30 for the fit between the observed and the calculated scattering curves (Fig. S6D-E). However, IgE contains five Asn-linked glycans attached to each epsilon chain that were not modelled and the FcεRIα ectodomain has seven Asn-linked glycans that were only modelled to a minor degree, and therefore a total 34 kDa of glycan had not been modelled. Addition of 17 full glycans and rigid body optimization of these glycans with the protein part fixed gave χ^2^ values of 1.9 (cryo-EM, Fig 3C), 2.7 (ns-EM major, Fig 3D) and 5.9 (ns-EM minor). Hence, the SAXS data suggested that the minor ns-EM conformation may not be relevant but failed to clearly distinguish between the cryo-EM and the major ns-EM conformation of the complex.

The IgE antibody is often depicted schematically as a quite bent molecule in order to align with fluorescence spectroscopy measurements that suggested distances of 71 Å and 75 Å between the antigen-binding site and the C-terminus of the IgE heavy chain in FcεRI bound and fluid phase IgE, respectively (10). In agreement with these measurements, there is 70-74 Å between IgE heavy chain Ser100 at the paratope of the proximal Fab and Ala542 at the C-terminal end of the same ε-chain in the models fitted to the cryo-EM 3D class and the major ns-EM 3D class (Fig. 3E). Together, our SAXS data and the prior spectroscopy data provide orthogonal evidence that two of the IgE conformations we observe by EM are relevant for the FcεRI bound state and not induced by the EM sample preparation methodology.

It is clearly of strong interest to examine how these IgE conformations translate to the intact tetrameric membrane embedded FcεRIαβγ_2_. Despite the lack of a detailed atomic structure of this receptor and the fact that we do not know exactly how the α-chain ectodomain is oriented spatially relative to the transmembrane part of the receptor, it appears that the two solution relevant conformations of the FcεRI bound IgE are likely to have different properties with respect to antigen recognition (Fig. 3F). If receptor bound IgE adopts the conformation observed by cryo-EM, the Fab arms and antigen binding site may be roughly parallel to the membrane. In contrast, if IgE adopts the major ns-EM conformation, one Fab arm is likely to be pointing away from the membrane and capable of reaching towards antigens located further from the surface of the FcεRI presenting cell.

### Increased Fab mobility lowers antigen stimulated IgE receptor activity

To understand why the IgE Fab-Fc hinge is rigid, we aligned sequences of the epsilon heavy chain (Fig. S7A). The hinge residues Ser222-Pro227 connecting the Cε1 domain in the Fab to the Fc Cε2 domain in the IgE Fc are not highly conserved, but a constant feature is a nonpolar residue (F/V/I/L) at the position equivalent of Phe225 in our structure. The resolution of our cryo-EM 3D reconstruction prevents us from performing a detailed analysis of the role of Phe225 in the hinge, but it is clear that this residue experiences different surroundings at the two Fab-Fc hinges. At the proximal Fab, Phe225 appears to be close to Tyr313 in the connected Cε2 domain and not in the proximity of the light chain (Fig. S7B). At the distal Fab, Phe225 is close to Phe318 in the Cε2 domain and the C-terminal residues of the light chain including the light chain Cys217 engaged in a disulfide bond with Cys131 in the heavy chain (Fig. S7C). Notably, the presence of aromatic residues at positions 313 and 318 is well conserved (Fig. S7A). For the conformations of FcεRI bound IgE observed by negative stain, we cannot define the linker conformations. But, as the overall locations of the proximal Fab arms are rather different from that observed by cryo-EM (Fig. 3A), there appears to be four different conformations of the linker for FcεRI bound IgE that supports a stable Fab-Fc association.

The EM studies of FcεRIα bound IgE presented here and our prior study of free IgE and IgE in complex with ligelizumab (15) provide convincing evidence that the Fab arms are associated with the nearby Cε2 domains in a limited number of stable conformations. In order to assess the importance of IgE rigidity in the context of its interplay with the receptors, we prepared the HMM5L variant of the HMM5 IgE with ten flexible residues inserted between Arg223 and Asp224 (Fig 4A). The flexibility of HMM5L was verified by ns-EM (Fig. S8A), but the binding kinetics determined for the HMM5 and HMM5L interaction with immobilized FcεRIα ectodomain were very similar (Fig. 8B-E). Hence, the stable association of the Fab arms with the Fc does not contribute to affinity for the receptor FcεRI.

**Figure 4.**
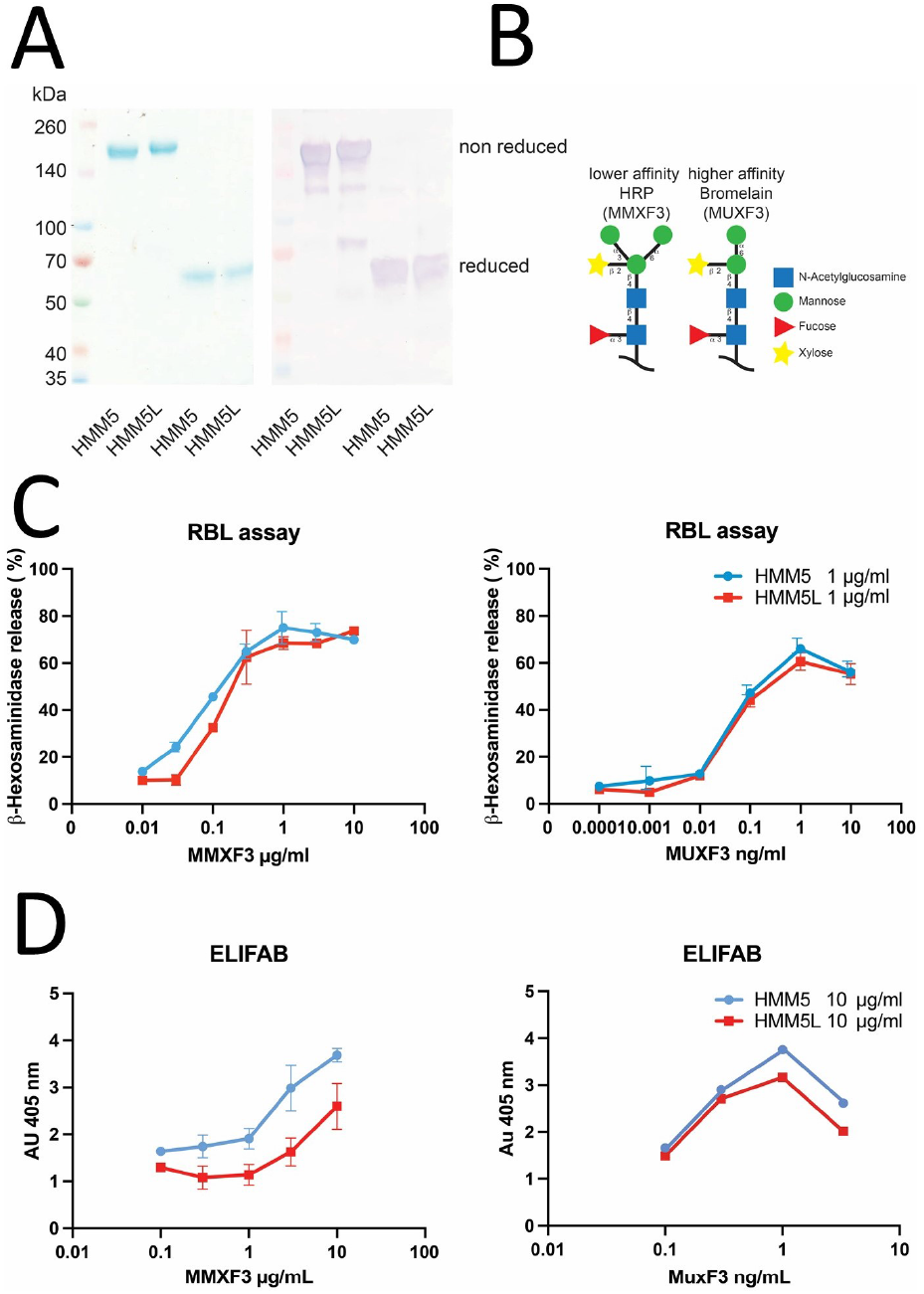
Introduction of flexibility in the hinge may alter the interaction of IgE with both its receptors. A) Right, SDS-PAGE analysis of HMM5 IgE and HMM5L IgE under non-reducing and reducing conditions after size exclusion chromatography. The band around 170 kDa represents the fully assembled antibody, whereas the band at 60 kDa under reducing corresponds to the ε-chain. The light chains are not visible under these conditions. Left, the identity of IgE was further confirmed by immunoblot using an anti-human IgE antibody labeled with alkaline phosphatase for detection. B) Schematic drawing of the two glycans recognized by the HMM5 IgE. The lower affinity antigen MMXF3 is found on horse radish peroxidase (HRP), the higher affinity antigen MUXF3 on pineapple stem bromelain. C) Binding and cross-linking of the high-affinity receptor FcεRI was assessed by detection of β-hexosaminidase release from RBL-S38X cells upon passive loading with IgE followed by exposure to the antigens. Both IgE variants elicit β-hexosaminidase release, but HMM5L is less active with the lower affinity MMXF3 antigen. D) The ability of IgE to form immune complexes and the subsequent binding to immobilized CD23 was analyzed in ELIFAB assays. A clear reduction of complex binding to CD23 for HMM5L compared to HMM5, in particular for the lower affinity antigen, suggests a more efficient complexation or sterically different complex architecture.

For the functional comparison of HMM5 and HMM5L, we took advantage of the fact that the HMM5 IgE exhibits reactivity to two similar carbohydrate structures, MUXF3 and MMXF3 differing by a mannosyl group (Fig. 4B), but with clearly different affinities (20). The potential of HMM5 IgE and HMM5L for activation and degranulation of effector cells by FcεRI cross-linking was assessed by measuring β-hexosaminidase release from RBL-SX38 cells, rat basophilic leukemia cells expressing the human FcεRI. Since receptor cross-linking by a monoclonal IgE like HMM5 demands the availability of more than one epitope per allergen we used horseradish peroxidase (HRP) carrying the MMXF3 glycan and a MUXF3 conjugate, consisting of bromelain-derived glycans coupled chemically to HSA, both of which display multiple cross-reactive carbohydrate determinant (CCD) epitopes. Both HMM5 and HMM5L exhibited clear β-hexosaminidase release as marker for cellular activation for both antigens, although as expected at different concentrations (Fig. 4C). However, for the lower affinity antigen MMXF3, a noticeable shift in the sensitivity of activation was observed. The calculated EC_50_ ratio (HMM5L/HMM5) for activation with MMXF3 was 1.4 ± 0.3 compared to 1.0 ± 0.2 for MUXF3, suggesting that the introduction of the linker compromised the activity of IgE in this assay for the lower affinity antigen MMXF3.

The low affinity IgE receptor CD23 is crucial for binding, uptake and presentation of allergens in form of allergen/IgE complexes and considered as another important driver of allergic immune responses. Therefore, our IgE variants were also evaluated for facilitated antigen binding (FAB) to CD23 using a surface-based variant of the FAB assay, the ELIFAB assay (24). HMM5 and HMM5L were used for formation of complexes with either the MUXF3 conjugate or MMXF3 (Fig. 4D). These complexes bound efficiently to CD23, corroborating the general capability of the HMM5 IgE to form immune complexes of higher order and facilitate allergen binding via CD23. Notably, the binding of the immune complexes to CD23 was clearly diminished for the HMM5L IgE compared to the parental HMM5 IgE, again in particular for the lower affinity MMXF3 antigen. This reduction in IgE/antigen complex binding might be based on a varying complexation efficacy despite equivalent affinity or, alternatively, sterically different complex architectures resulting in a change in binding behavior. Taken together, data obtained with the two different functional assays suggest that rigidity of the Fab-Cε2 hinge, depending on the antigen and its architecture, is important for the biological function of IgE.

## Discussion

Here we have provided firm structural evidence that IgE can adopt distinct defined conformations when bound to the ectodomain of FcεRIα, but the relevance of the minor ns-EM conformation remains uncertain. Also, we cannot exclude that the IgE conformation when bound to cellular FcεRI differs from those of the IgE-FcεRIα complex capture by EM, but since IgE is well separated from the membrane and the transmembrane parts of FcεRI, the cryo-EM IgE and possibly also the major ns-EM conformations we observe are likely to represent *in vivo* relevant conformations. The energetic landscape of proteins may differ substantially between bulk solution and the air-water-interface in cryo-EM (25). Varying degrees of distortions have also been observed in room temperature EM where the sample is stained with a heavy metal salt and adsorbed to a carbon surface (26). These distinct environments potentially cause the difference in conformational state that we observe between cryo- and ns-EM, sample preparation appears to stabilize specific states that in bulk solution would be in dynamic equilibrium. To ensure that the conformations in our final structures were not significantly biased by the initial particle picking strategy, we compared template matching with references based on the ns-EM and cryo-EM structures, but did not observe any differences in the picked particles. Furthermore, as traditional classification in cryoEM is dependent upon discrete differences existing between states that can be classified out, we also used cryoDRGN to investigate whether there is a large degree of continuous flexibility in the data (27). However, we did not observe any large continuous flexibility except between the constant and variables domains in the Fab.

Fluorescence spectroscopy experiments suggested a distance of 71 Å and 75 Å between the antigen-binding site and the C-terminus of the IgE heavy chain in FcεRI bound and fluid phase unbound IgE, respectively (10). These two IgE conformations observed by EM are compatible with these distance for one Fab arm. It should be noted that in FRET spectroscopy, distances beyond 10 nm are not well resolved and the longer distance between the distal Fab paratope and IgE C-terminus may be difficult to measure with this technique. With respect to the geometry of Fab arms, titration of an anti-DNP IgE with bivalent antigens suggested a separation of 13 nm between the two DNP paratopes (28) which is compatible with the minor ns-EM conformation. In the cryo-EM conformation, the distance is 17 nm, but divalent ligands capable of spanning this distance were not evaluated (28).

Since the IgE Fc and IgE bind FcεRIα with very similar affinity (29, 30) and also shown by us here (Fig. S8B-E), we do not expect significant differences between the observed IgE conformations in affinity for this receptor. With respect to the second receptor CD23, the binding site is located in the Cε3 domain of IgE (31), is distinct from the FcεRIα binding site, and requires the closed conformation within the Cε3-Cε4 part of IgE Fc. We notice that binding of CD23 to IgE appears to be incompatible with the observed IgE conformations since the distal Fab is predicted to exert steric hindrance on CD23. Further biophysical and ultrastructural characterization of IgE bound to FcεRI and CD23 presenting cells is required to clarify the role of the observed IgE conformations and how they may change in response to antigen binding and receptor crosslinking.

Like IgE, IgM has a short hinge between the Fab and the Fc and four constant domains Cμ1-Cμ4 in the heavy chain. In a recent cryo-EM structure of pentameric IgM, one of the Cμ2 domain tandems and the associated Fab arms were observed in a defined position (32). The angle between IgM Fab arms is ∼28° and a similar angle was also observed in a tomography study of IgM in complex with complement component C1 (33). Likewise, a recent structure of B cell receptor IgM associated with the CD79αβ heterodimer features a similar orientation of the two Fab arms (34). Hence, even though IgE and IgM both have short hinge regions and their Fab arms stably attached to the Cε2 and Cμ2 domains, respectively, the IgE conformations presented here are clearly distinct from the Fab-Cμ2 arrangement observed in multiple structures of IgM (Fig. S7D).

Our functional assays argue that structurally defined hinge conformations play a significant role in the biological activity of IgE. We also notice that a recent preprint reports on experiments in which polyvalent DNP-BSA was used as antigen. Similar to our own assays in Figure 4C, a reduction in degranulation of RBL cells was observed when a flexible linker was introduced between the conjugated DNP molecules and the BSA carrier (35). Hence, introduction of flexibility in the antigen and the IgE linker region both reduce the biological activity of the antigen-IgE immune complexes. It is well described that different antibody isotypes exhibit varying capability to form complexes of varying size and architecture with the antigen in solution (13). Although the use of a monoclonal IgE simplifies the analysis in the functional assays performed in this study, the accompanying use of a polyvalent antigen for cellular activation and complex formation makes an extrapolation somewhat difficult. With more than 1000 allergens approved by the WHO/IUIS, individual target structures probably exhibit different susceptibility to rigid or flexible IgE molecules. Nevertheless, it appears reasonable that in the polyvalent setting of this study, an impact is primarily seen for a lower affinity interaction. Inversely, a high affinity interaction combined with increased avidity might be less amenable to reflecting the changes imposed by hinge flexibility in the biological activity of IgE. Similarly, allergens of different three-dimensional structure and size might preferentially interact with the one or the other IgE conformation. Epitopes in direct proximity seem to establish more potent allergens, a concept which is supported by the fact that a large portion of allergens is actually rather small. Most of these studies however were based on single IgE antibodies with engineered or synthetic epitopes. In an allergic patient with the presence of an IgE repertoire of varying complexity, the impact of molecular rigidity, the conformational state and the consequences for the interaction with the two receptors could easily be profoundly stronger.

## Author contributions

RKJ conducted all steps of the cryo-EM analyses and prepared the FcεRIα ectodomain. MM expressed and purified IgE and conducted the functional assays. RG performed all the experimental work related to SAXS data. MHJ collected and analyzed ns-EM data. EZ and GRA supervised research, analyzed data and wrote the paper together with RKJ, RG, MHJ and MM.

## Acknowledgements

We are grateful to assistance with SAXS data collection offered by the EMBL Hamburg staff. We acknowledge access to the cryo-EM facilities at the UK’s national Electron Bio-imaging Centre (eBIC), and to Mirian Weckener and Dan Clare at eBIC for their assistance with preparing grids. We also thank the EM facility at Aarhus University for their assistance with data collection and Jesper L Karlsen for help with EM data processing.

## Materials and Methods

Expression of HMM5 and HMM5L IgE was essentially performed as described (36). To create the HMM5L variant with a flexible hinge, a 10 amino acid (SG_4_)_2_ linker was inserted between residues 223 and 224 of the HMM5 IgE. The ectodomain of the FcεRIα subunit (residues 26-205) was prepared essentially as described in (37). The ELIFAB assay was conducted according to (38). In the bio-layer interferometry experiments, association of fluid phase HMM5, HMM5L or IgE Fc with immobilized biotinylated FcεRIα was measured. Cryo-EM data were collected at the eBIC facility while ns-EM data were acquired at the Aarhus University iNANO EM facility. EM data were processed in CryoSPARC (39) and the resulting 3D reconstructions were inspected and models fitted in Chimera-X and COOT (40, 41). Refinement and validation was done with PHENIX (42). SAXS data analysis and rigid body refinement was carried out with ATSAS (43). Full methods are presented in the supporting material. Figures were prepared in Pymol (https://pymol.org/pymol.html) or Chimera-X.

