## Supporting information for "The dynamics of hinge flexibility in receptor bound immunoglobulin E revealed by electron microscopy"

### Methods

#### Protein expression and purification.

Expression of HMM5 IgE was essentially performed as described (1). Briefly, Sf9 cells were cultivated in HyClone Sfx-insect medium (Cytiva). Baculovirus production was performed by using the Bac-to-Bac system following the manufacturer's recommendations. HMM5 and HMM5L plasmids were transformed into DH10Bac™ *E.coli* and the recombinant bacmid DNA was isolated and transfected into SF9 cells using Cellfectin II (Thermo Fisher Scientific). After two rounds of virus amplification, the appropriate virus concentration for expression was analyzed by infecting Sf9 cells with different amounts of virus and analyze protein expression by immunoblot 5 days after infection. For expression 500 ml Sf9 cells $1x10^6$ cells /ml were infected with virus and incubated under shaking for 5 days at 27°C.

To create the HMM5L variant with a flexible hinge, a 10 amino acid $(SG_4)_2$ linker was inserted between amino acid residues 223 and 224 of the HMM5 IgE. In order to insert the SG linker, a gene string (GenArt) containing the IgE-Fc sequence fused to the SG linker was ordered and cloned between *SalI* and the *BglII* of IgE-Fc. The IgE-FC linker was then inserted via *XhoI* and *HindIII* into pFastBac dual containing HMM5 kappa light chain and the heavy chain variable region (pFBM5hIgE). After expression, the antibodies were purified from the culture medium using a capture with a 5 ml HisTrap excel column (Cytiva). The column was washed with 20 column volumes (CVs) 20 mM NaPO4, 0.5 M NaCl, 30 mM imidazole, pH 7.4 and eluted with 15 CVs 20 mM NaPO4, 0.5 M NaCl, 500 mM imidazole pH 7.4. The concentrated eluate was further purified by size exclusion chromatography using on a HILoad 16/600 Superdex 200pg column (Cytiva) equilibrated in 20 mM HEPES, 50 mM NaCl pH 7.0. A final SEC chromatography step was performed on a 24 ml Superdex 200 increase (Cytiva) equilibrated in 20 mM HEPES pH 7.5, 150 mM NaCl.

For expression in HEK293 cells, the cells were incubated in DMEM high glucose L-glutamine (Sigma-Aldrich) supplemented with 10% fetal calf serum and 0.1 kIU/L penicillin and 0.1 mg/mL streptomycin. For transfection, $5x10^5$ cells per 6-well were seeded and 20 h later transfected with 4 µg of DNA using polyethylene imine (Polysciences) in a DNA:PEI ratio of 1:3. For stable expression, the cells were passaged in a ratio 1:5 3 days after transfection and exposed to zeocin (invivogen). After 2 weeks of selection, single colonies were isolated and protein expression was analyzed by immunoblot. For expression, cells of a confluent T175 flask were transferred to an 850 $cm^2$ rolling bottle with 500 ml DMEM medium and incubated rolling for 14 days.

The ectodomain of the FcεRIα subunit (residues 26-205) was expressed in HEK293S cells transfected with PEI as described in (2). The cell supernatant was collected after 4 days and applied to a 5 mL HisTrap Excel (Cytiva). The column was washed twice with 25 mL of 20 mM TRIS pH 8.0, 500 mM NaCl, and FcεRI was eluted in 20 mL of 20 mM TRIS pH 8.0, 250 mM Imidazole. The FcεRIα was added 1:100 (w:w) TEV protease and was dialyzed 2 hours against 2 L of 20 mM HEPES pH 7.5, 150 mM NaCl at room temperature followed by 16 hours at 4 °C against 2 L of 20 mM HEPES pH 7.5, 150 mM NaCl. The protein was applied to a 5 mL HisTrap HT, and FcεRIα was collected in the flow through of the column. The protein was concentrated using a vivaspin 6 (Merck) with a molecular weight cut off of 10 kDa, and thereafter applied to a 24 ml Superdex 200 increase (Cytiva) equilibrated in 20 mM HEPES pH 7.5, 150 mM NaCl. The IgE-FcεRIα complex was formed by mixing IgE with a two-fold molar excess of the receptor. The complex was incubated for at least 30 minutes on ice, before being applied to a 24ml Superdex 200 increase SEC column (Cytiva) equilibrated in 20 mM HEPES pH 7.5, 150 mM NaCl.

#### Mediator release assay.

RBL-SX38 cells were cultivated in a $CO_2$ incubator at 37°C with MEM, 15% heat-inactivated fetal calf serum with 0.1 kIU/L penicillin, 0.1 mg/mL streptomycin, 1 mM sodium pyruvate and 1 mg/mL G418 (Sigma-Aldrich). In vitro degranulation was analyzed as described previously (3) (4). One day prior to the assay, $1x10^5$ cells per 96 well were seeded. After sensitization with 100 ng of IgE in culture-medium for 2h, different concentrations of CCD-carrying proteins were added in Tyrode's buffer (10mM HEPES,13 mM NaCl, 5mM KCl, 1.4mM $CaCl_2$, 1.0 mM $MgCl_2$, pH 7.4, 5.6 mM glucose, 0.1% w/w BSA) and incubated for 60 min at 37 °C. As a reference, cross-linking was achieved by 100 ng polyclonal rabbit anti-human IgE antibody (Dako A0094). β-Hexosaminidase release of viable versus lysed cells was assessed with 8 mM p-nitrophenyl N-acetyl-glucosaminopyranoside in 0.1 M citrate buffer pH 4.5. The reaction was stopped with 0.1 M carbonate buffer pH 10 and the absorbance measured at 405 nm.

#### CD23 binding in the ELIFAB assay.

In order to analyze the immunocomplex formation and their binding to CD23, a surface-based ELIFAB assay was performed (5). Immunocomplexes were formed by adding 20 µL HMM5 or HMM5L IgE (10 µg/ml in RPMI) to 20 µL RPMI and 5 µL antigen followed by incubation for 1h at 37°C. The preincubated immunocomplexes were transferred to overnight coated and BSA-blocked recombinant CD23 (625 ng/ 96 well in PBS (R&D systems 123-FE-050)). After washing; IgE antigen complexes bound to immobilized CD23 were detected by adding biotin-conjugated anti-human IgE antibody (0.5 µg/mL) (BD-Pharmingen Cat 55585) followed by detection with ExtrAvidin AP (Sigma E2636) and p-nitrophenyl phosphate 5 mg/mL (Sigma 4876) in alkaline phosphate buffer.

#### Bio-layer interferometry

All BLI experiments were performed on an Octet Red96 (Sartorius) at 30°C and shaking at 1000 rounds per minute using streptavidin biosensors (Sartorius). The experiments were performed in 20 mM HEPES pH 7.5, 150 mM NaCl, and regeneration was performed in three rounds of 30 seconds using 2 M NaCl, 100 mM Acetate pH 4.0. First FcεRIα was biotinylated O/N at 4°C in 20 mM HEPES pH 8.0, 150 mM NaCl, 10 mM ATP, 50 µM biotin, 10 mM $MgCl_2$ and BirA in 1:10,000 (w:w). BirA was removed by using a 24 ml Superdex 75 increase (Cytiva) SEC column. The streptavidin sensors were loaded with 5 µg/mL biotinylated FcεRIα for 2 minutes, and subsequently washed thrice with running buffer. Association was then measured to IgE Fc (12.5, 6.3, 3.1, 1.6, 0.78, 0.39 nM), HMM5 (25, 12.5, 6.3, 3.1, 1.6, 0.78, 0.39 nM) or HMM5L (50, 25, 12.5, 6.3, 3.1, 1.6, 0.78n nM) for 300 seconds, followed by a 300 seconds dissociation period. The association was modeled as $R(t)=R_{max}([IgE]/([IgE]+K_D)(1-exp(-t(k_{on}[IgE]-k_{off})))$, $K_d=k_{on}/k_{off}$, and the dissociation was modelled as a first order exponential decay, $R(t) = R(300)exp(-k_{off}(t-300\ s))$. The given values correspond to the average of three (HMM5 IgE, IgE Fc) or two (HMM5L IgE) experiments ± the standard deviation.

#### Cryo-EM grid preparation, data collection and processing

The purified IgE-FcεRIα complex was concentrated to 3.2 mg/mL, and applied to Quantifoil Active grids (SPT Labtech) using the Chameleon (SPT Labtech) at the United Kingdom national electron Bio-Imaging Centre (eBIC). Micrographs were collected on a 300 kV Titan Krios microscope (ThermoFischer) at eBIC, equipped with a Bio-quantum-K3 detector and an energy filter (Gatan). For data collection spot size 5 and a C2 aperture of 70 µm were used, while micrographs were collected with EPU using at 105k x magnification, and a physical pixel size of 0.829 Å. The data were collected in super resolution mode with an assumed defocus of -1.8 µm to -0.8 µm, varied in 0.2 µm steps. Each micrograph was collected with a dose of 59.16 $e^-/Å^2$ divided over 40 frames. Raw super-resolution movie frames were imported into CryoSparc v3 (Structura Biotechnology 33 Inc) (6). The movies were initially aligned using patch motion correction with a maximum alignment resolution of 5 Å, B factor blurring of 500 Å, and cropping the output two-fold to the physical pixel size. The CTF was estimated using patch CTF estimation with default parameters. Movies with defocus outside of 0.5 µm and 4 µm were removed, corresponding to 204 movies. Particles were picked using the blob picker giving a total of 2,936,131 particles which were extracted with 5-fold binned data. Iterative rounds of 2D classification yielded a dataset corresponding to 634,649 particles. The particles were cleaned with a round of ab initio model generation using three classes. The two best classes were refined using 3D heterogeneous refinement with three classes. The best two classes from heterogeneous refinement, corresponding to 573,328 particles were re-extracted at 2-fold binning and a consensus model was refined. The consensus model was subsequently refined using non-uniform refinement (7). The refined model was used to perform local motion correction, followed by a new round of non-uniform refinement. The resulting angles were used to perform focused refinement on the FcεRI and IgE Cε2-4 to obtain well refined angles for the least flexible part of the structure. Subsequently, we performed local CTF alignment, and a final round of non-uniform refinement giving corresponding to the final cryo EM map of the full IgE at a global resolution of 4.1 Å. A final round of focused refinement on FcεRI and IgE Cε2-4 gave the final 3D reconstruction for focused refinement with a resolution of 3.8 Å.

#### Structure refinement and analysis

Chimera-X (8) was used for map inspection and initial fitting of models to 3D reconstructions. A model of the Fab containing the Cε1 domain was prepared with alphafold2 (9). Programs from the PHENIX package (10) version 1.19 or 1.20 was used for real space refinement, analysis and validation of cryo-EM structures. The graphics program COOT (11) was used for manual rebuilding of models prior to real space refinement in PHENIX. All figures presenting atomic models and 3D reconstructions were prepared with PYMOL, https://pymol.org/pymol.html or Chimera-X. To establish the optimal conformation of the two Fab moieties, models in a library containing 2468 unique Fab structures was superimposed onto the constant domain in the two light chains of an initial model refined against the consensus map. The correlation between the two fitted Fab molecules and the map was calculated with Chimera-X. The Fab in pdb entry 4dw2 was found to have the highest correlation for both Fab arms. An optimal model for the full IgE was next constructed by superposition of the variable domains from the structure of the HMM5 Fab in pdb entry 5i8o onto the variable domains of the two models of 4dw2 fitted to the two Fab arms in the model of the IgE-FcεRIα complex.

#### Negative stain EM

A carbon-evaporated copper grid (G400-C3, Gilder), was glow-discharged at 25 mA for 45 seconds using an easiGlow instrument (PELCO). The IgE-FcεRIα complex was applied on the grid in 3 µL at 3 µg/mL and incubated for 2 min. The grid was then washed in 3 µL deionized water twice, washed in 3 µL Nano-W (Nanoprobes) and, finally, incubated in 3 µL Nano-W for 90 s. The stain was blotted away and the grid was air-dried before being imaged on a 120 kV Tecnai G2 spirit electron microscope. Automated data collection was performed using the SerialEM (12), at a nominal magnification of 67,000x and -1 µm defocus. 294 micrographs were collected and processed in cryoSPARC (6). The two rigid bodies derived from the IgE-FcεRIα complex were fitted into the 3D reconstructions with Chimera-X and map-model correlations were calculated at 25 Å resolution.

#### SAXS analysis

SEC-SAXS measurements were performed on the IgE-FcεRIα complex at P12 beamline at Petra III, DESY (Hamburg, Germany). The HMM5 IgE and the purified FcεRIα ectodomain were dialyzed against 2 L of 25 mM HEPES pH 7.5, 150 mM NaCl for 24 h at 4°C and concentrated to 5.5 mg/ml. To reconstitute the IgE-FcεRIα complex, IgE was mixed with a two-fold molar excess of FcεRIα. The protein solutions were centrifuged at 15000 rpm for 15 min at 4°C prior to injection onto a Superdex 200 Increase 5/150 column (Cytiva) equilibrated in 25 mM HEPES pH 7.5, 150 mM NaCl for the in-line SEC-SAXS measurements. Programs in the ATSAS package version 3.0.3 or 3.0.5 (13) were used for data analysis using experimental data with a maximum q value of 0.2 $Å^{-1}$. Data analysis and reduction was performed with CHROMIXS and PRIMUS. Atomic models were compared with the experimental data using CRYSOL. A model of the fully glycosylated IgE-FcεRIα in the cryo-EM conformation was created by addition to each relevant Asn of the NAG-NAG-MAN moiety in Coot and subsequent completion of the glycan by superposition of the mannose residue with the corresponding mannose in the glycan in pdb entry 3ry6. Models of the glycosylated ns-EM models were prepared by superposition of the Fab and associated glycans from the glycosylated EM model. Rigid body optimization of the glycans was performed with CORAL using a 6 Å distance restraint between the relevant glycan and the closest NAG residue. The protein parts of the models and the glycans attached to Asn391 in the Fc were kept constant during rigid refinement.

*Table S1. Statistics for cryo-EM data collection, refinement and validation*. The lower part of the table was prepared with phenix.validation_cryoem.

| **Data collection** | | | |
|---|---|---|---|
| Microscope | FEI Titan Krios | No. of frames | 40 |
| Magnification | 105,000 | Defocus range (µm) | 0.8-1.8 |
| Voltage (kV) | 300 | Pixel size (Å) | 0.829 |
| Electron exposure ($e^{-}/Å^2$) | 59.16 | Symmetry imposed | C1 |
| | | No. of micrographs | 11015 |
| **Deposited data** | **Focused** | **Consensus** | |
| PDB/EMDB entry | 8C1B/ EMD-16377 | 8C1C/EMD-16378 | |
| Atoms | 12675 (Hydrogens: 6106) | 19267 (Hydrogens: 9284) | |
| Residues | Protein: 807 | Protein: 1237 | |
| Bonds (RMSD) | | | |
| Length (Å) (# > 4sigma) | 0.007 (0) | 0.007 (0) | |
| Angles (°) (# > 4sigma) | 1.257 (16) | 1.156 (14) | |
| MolProbity score | 1.42 | 1.23 | |
| Clash score | 3.18 | 2.40 | |
| Ramachandran plot (%) | | | |
| Outliers/ Allowed/ Favoured | 0.25/4.24/95.51 | 0.16/3.18/96.66 | |
| Rotamer outliers (%) | 0.28 | 0.00 | |
| ADP (B-factors) | 6569 | 9983 | |
| Protein (min/max/mean) | 74.57/252.79/131.77 | 33.87/999.99/346.65 | |
| Ligand (min/max/mean) | 77.58/204.62/119.18 | 114.39/256.02/176.60 | |
| Data | | | |
| Box Lengths (Å) | 96.37, 122.28, 103.62 | 116.06, 122.28, 138.86 | |
| Supplied Resolution (Å) | 3.8 | 4.1 | |
| Resolution Estimates (Å) | Masked/Unmasked | Masked/Unmasked | |
| d FSC (half maps; 0.143) | 3.8/3.9 | 4.2/4.2 | |
| d model | 3.9/4.0 | 4.0/4.1 | |
| d FSC model (0/0.143/0.5) | 3.6/3.8/4.5<br>3.7/3.9/6.9 | 3.9/4.1/6.3<br>4.0/4.2/7.1 | |
| Map min/max/mean | -3.87/5.25/0.17 | -1.18/3.26/0.13 | |
| Model vs. Data | | | |
| CC (mask/box/peaks/volume) | 0.70/0.76/0.53/0.70 | 0.71/0.87/0.66/0.70 | |
| Mean CC for ligands | 0.74 | 0.71 | |

#### Legends to supporting information figures

Figure S1. Cryo-EM data processing. A) Workflow for cryoEM processing B) Orientation analysis. The equivalents of the rotation and tilt angle using the RELION convention were plotted against each other using a Mollweide projection. The number of particles in each bin is indicated by color. C) Fourier shell correlation of the consensus map for the FcεRIα-IgE complex. D) Fourier shell correlation for the focused refinement containing the FcεRIα-IgE Fc part. E) The two 3D reconstructions colored according to their local resolution. To the right, slices presenting the central parts of the maps are shown. Panel A and E were prepared with ChimeraX.

Figure S2. Details of the cryo-EM map and the model of the IgE Fc. A-D) Examples of map quality from the focused refinement in Cε2 (A), Cε3 (B), Cε4 (C), and FcεRIα (D). The map is contoured at 12σ. In panels A-E, carbon atoms in one IgE heavy chain are colored blue, while carbons in the other heavy chain are green. In panel D, carbon atoms in the receptor α-subunit are colored grey, notice the clear density for the glycan linked to Asn42 of the receptor. E) The Cε2 domains with the consensus cryo-EM map contoured at 6σ. Extra density is visible around Asn262 for both heavy chains in accordance with an Asn-linked glycan. Furthermore, unambiguous density is present for residues 275-284 as part of the lower β-sheet that contradicts the previously reported strand shift upon IgE Fc receptor binding (14). F) Comparison of the Cε2 domain from the IgE-FcεRIα complex with the crystal structure of Fc-FcεRIα (left, pdb entry 2y7q) and unbound Fc (right, pdb entry 5mol).

Figure S3. Examples of the cryo-EM consensus map and the model for both Fabs. A) The consensus map around the Cε1-hinge-Cε2 region for pFab (left) and dFab (right). B) Presentation of the entire Fabs in the consensus map with the three glycosylated asparagine residues in the Cε1 domains shown as orange spheres. C) Close-up on the Cε1 domains contoured at 6σ. Significant additional density is present around Asn165 in both Fabs in agreement with its position. This further supports the selected option 1 for the Fab orientation presented in figure 2A-B.

Figure S4. Details of the IgE-FcεRIα complex structure. A) Predicted alphafold2 model of the HMM5 IgE Fab with a Cε1 domain. Orange spheres mark the three asparagines with N-linked glycans (left). Notice the location of the Cε1 specific disulfide bridge Cys131-221 next to the inter-subunit disulfide C130-C217CL (right). B) Details of the interaction between the receptor and the two Cε3 domains. Left, the Cε3A site interacting with an aromatic cluster in the FcεRI α1 domain. Right, detailed view of the Cε3B subsite.

Figure S5. Negative stain EM of the IgE-FcεRIα complex. A) Workflow for negative stain processing. As all micrographs were taken close to -1 µm defocus, no CTF correction was performed. Particles partitioned into the 3 classes of the final heterogeneous refinement in a distribution of 36 %, 32 % and 31 %. The first and the last classes were very similar and were combined in a single refinement, giving rise to the major conformation, while the middle class showed a different angle between the F(ab)s. B) Fourier-shell correlation for the major conformation volume. C) Comparison of the ns-EM maps of the major conformation (grey) and the minor conformation (yellow). D) Docking of the cryo-EM model after removal of the proximal Fab demonstrates a significant better fit of the model to one of the two possible hands for both the ns-EM maps.

Figure S6. Small angle X-ray scattering analysis of the IgE-FcεRIα complex. A) SEC-SAXS profile obtained with the IgE:FcεRIα complex. B-C) Indirect Fourier transformation of the merged data and the pair distance distribution obtained with the ATSAS program GNOM. D-F) Theoretical scattering curves (black) calculated with CRYSOL based on the atomic models of the complex prior to completion of the Asn-linked glycans compared to the experimental data (blue). These fits can be compared to those in Figure 3B and 3D showing the fits for fully glycosylated models.

Figure S7. The conformations of the hinge regions are different. A) Conservation of the hinge and the hinge contact region in the Cε2 domain. B) Close-up on the hinge at the proximal Fab with Phe225 in contact with the Cε2B domain. C) At the distal Fab, the hinge regions is sandwiched between the Cε2A domain and the C-terminal end of the CL domain. D) Comparison of three Fab conformations of IgE with one of multiple similar Fab conformations from IgM (15). The IgE Fab-Cε2 conformations are clearly distinct from the IgM Fab-Cμ2 conformation.

Figure S8. Introduction of a flexible linker does not change the affinity for FcεRIα. A) Most populated 2D class averages from ns-EM analysis of HMM5L reveals that in this IgE variant the Fab arms are flexible relative to the Fc part of the antibody. These 2D classes can be directly compared to the 2D classes of the parental HMM5 IgE presented in reference (16). B-D) Bio-layer interferometry data for fluid phase IgE-Fc (panel B), HMM5 (panel C), and HMM5L (panel D) binding to immobilized FcεRIα ectodomain. The raw curves are shown in black and the fitted model is shown in red. E) The dissociation and rate constants for the analyzed FcεRIα-IgE complexes. The average values and their standard deviation are derived from three (HMM5, IgE Fc) or two (HMM5L) experiments.

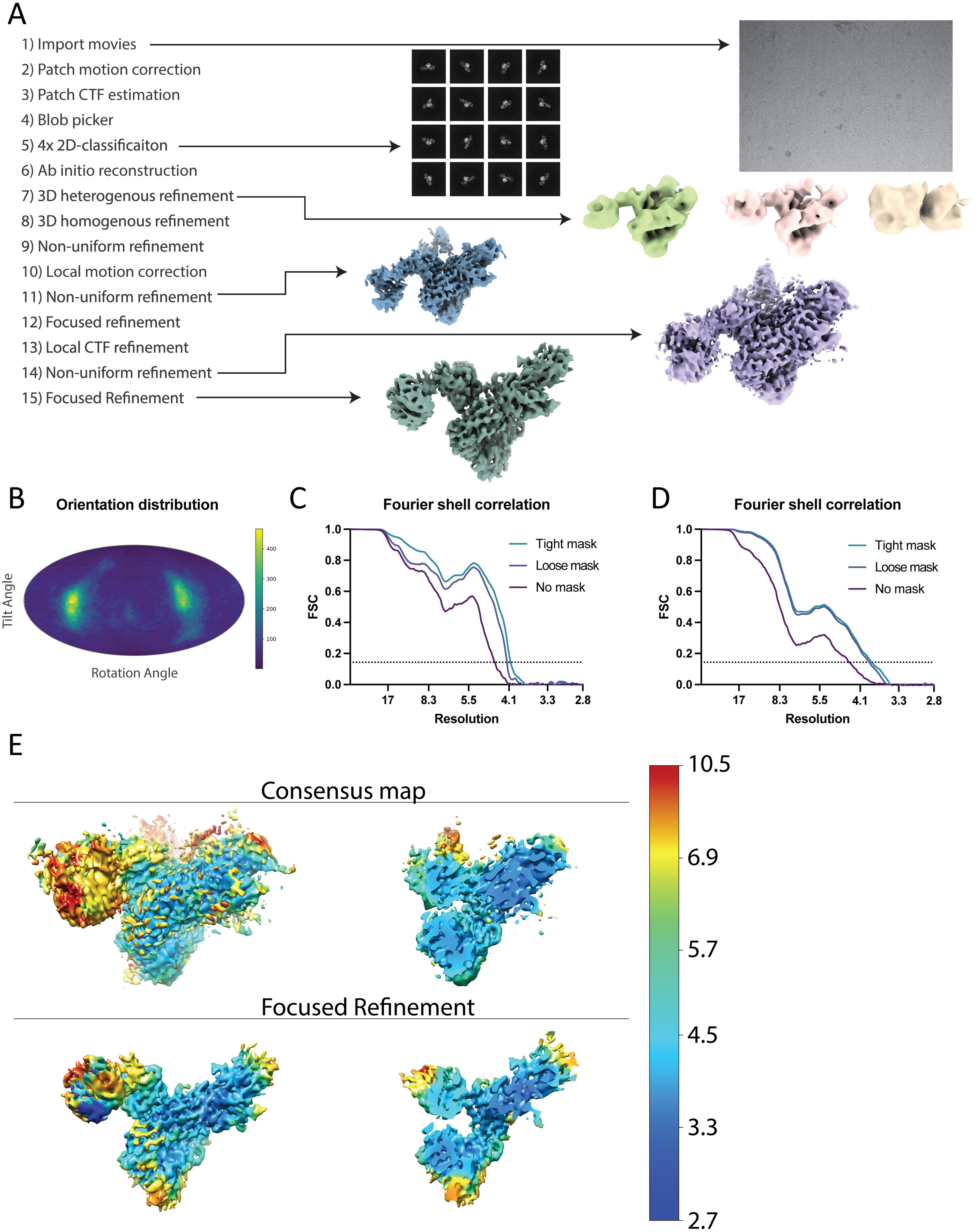
A
1) Import movies
2) Patch motion correction
3) Patch CTF estimation
4) Blob picker
5) 4x 2D-classificaiton
6) Ab initio reconstruction
7) 3D heterogenous refinement
8) 3D homogenous refinement
9) Non-uniform refinement
10) Local motion correction
11) Non-uniform refinement
12) Focused refinement
13) Local CTF refinement
14) Non-uniform refinement
15) Focused Refinement
B
Orientation distribution
Tilt Angle
Rotation Angle
400
300
200
100
C
Fourier shell correlation
FSC
Resolution
Tight mask
Loose mask
No mask
1.0
0.8
0.6
0.4
0.2
0.0
17
8.3
5.5
4.1
3.3
2.8
D
Fourier shell correlation
FSC
Resolution
Tight mask
Loose mask
No mask
E
Consensus map
Focused Refinement
10.5
6.9
5.7
4.5
3.3
2.7

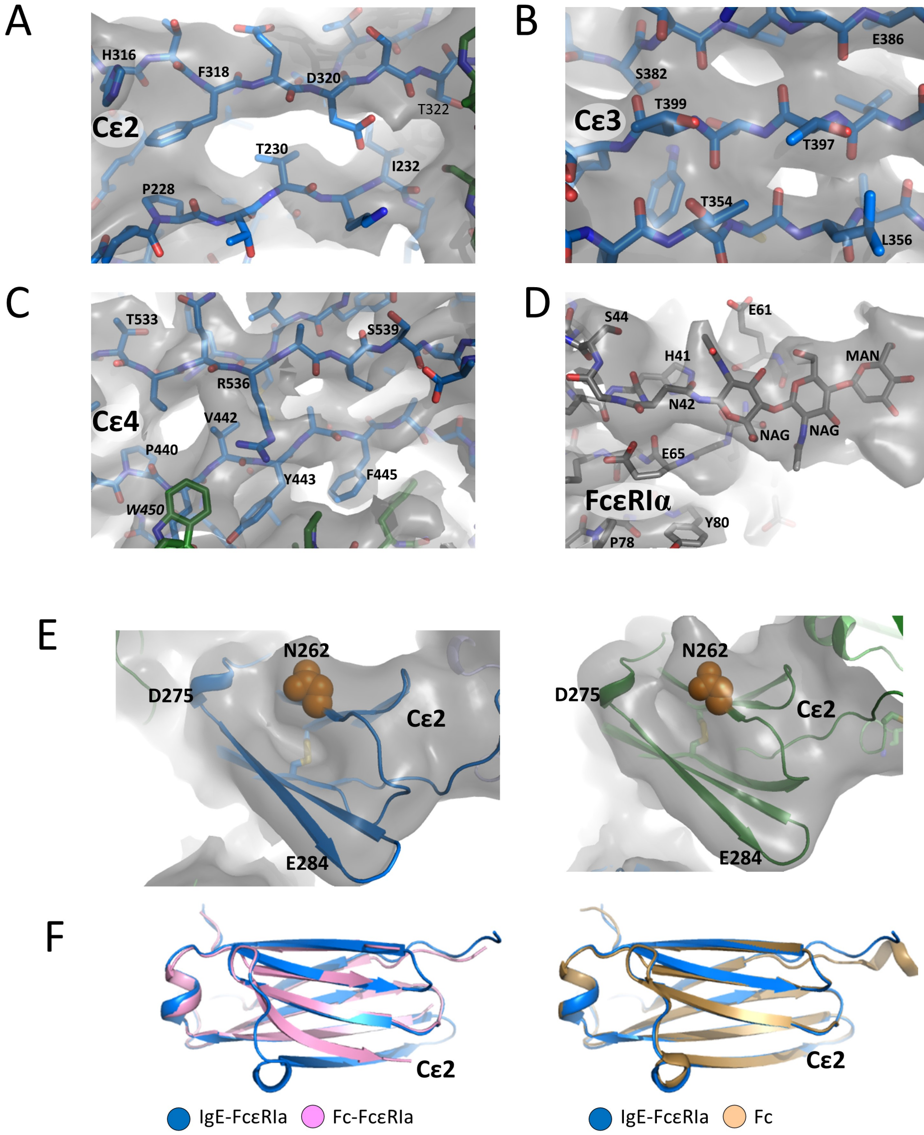

A
H316
F318
D320
T322
Cε2
T230
I232
P228
B
E386
S382
T399
Cε3
T397
T354
L356
C
T533
S539
R536
Cε4
V442
P440
Y443
F445
W450
D
S44
E61
H41
MAN
N42
NAG
NAG
E65
FcεRIα
Y80
P78
E
N262
D275
Cε2
E284
N262
D275
Cε2
E284
F
Cε2
IgE-FcεRIa
Fc-FcεRIa
Cε2
IgE-FcεRIa
Fc

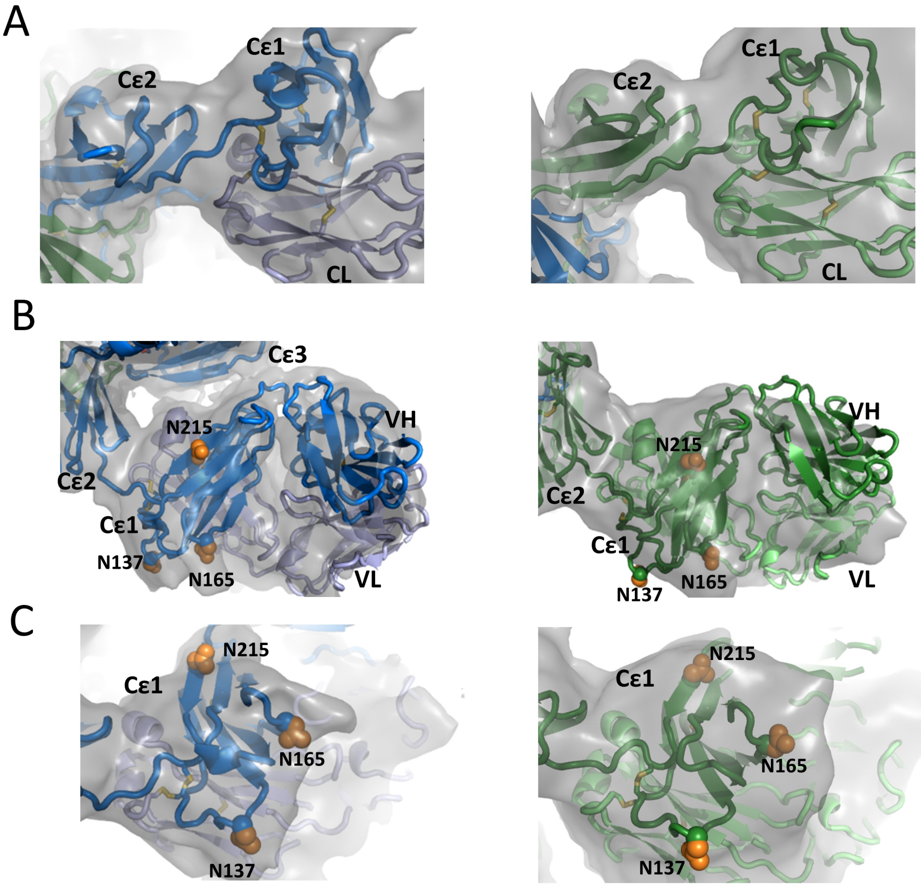
A
Cε2
Cε1
CL
Cε2
Cε1
CL
B
Cε3
N215
VH
Cε2
Cε1
N137
N165
VL
N215
VH
Cε2
Cε1
N137
N165
VL
C
N215
Cε1
N165
N137
Cε1
N215
N165
N137

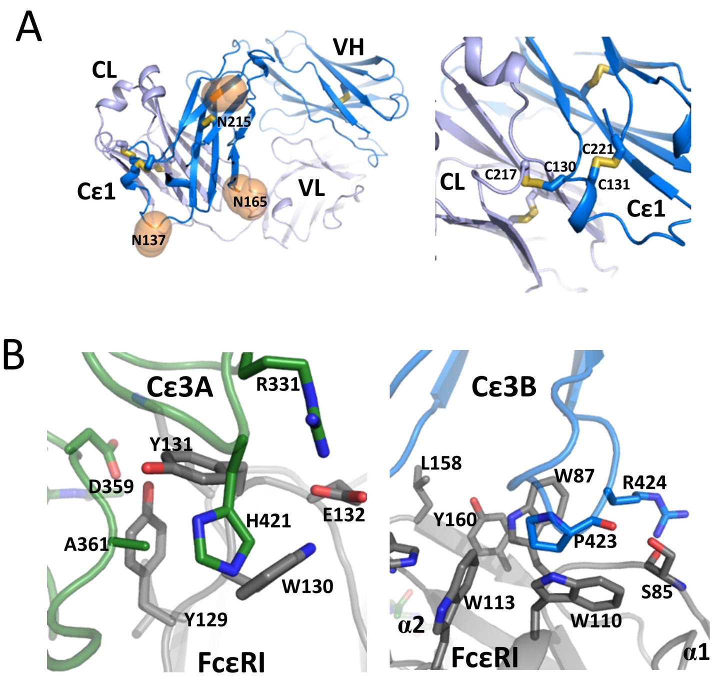
A
CL
VH
N215
Cε1
N165
VL
N137
CL
C217
C130
C221
C131
Cε1
B
Cε3A
R331
Y131
D359
H421
E132
A361
W130
Y129
FcεRI
Cε3B
L158
W87
R424
Y160
P423
S85
W113
W110
α2
FcεRI
α1

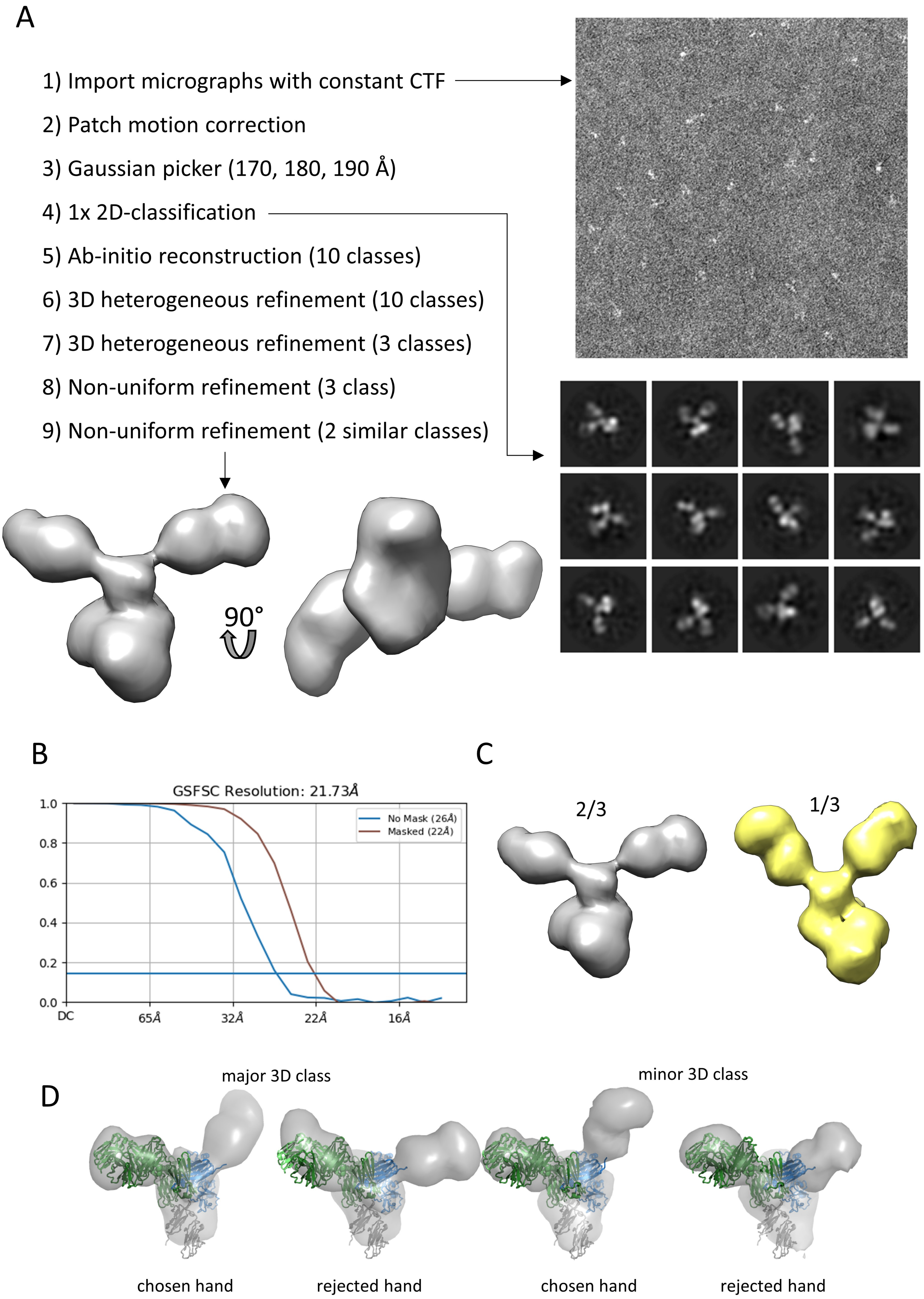
A
1) Import micrographs with constant CTF
2) Patch motion correction
3) Gaussian picker (170, 180, 190 Å)
4) 1x 2D-classification
5) Ab-initio reconstruction (10 classes)
6) 3D heterogeneous refinement (10 classes)
7) 3D heterogeneous refinement (3 classes)
8) Non-uniform refinement (3 class)
9) Non-uniform refinement (2 similar classes)
90°
B
GSFSC Resolution: 21.73Å
No Mask (26Å)
Masked (22Å)
1.0
0.8
0.6
0.4
0.2
0.0
DC
65Å
32Å
22Å
16Å
C
2/3
1/3
D
major 3D class
minor 3D class
chosen hand
rejected hand
chosen hand
rejected hand

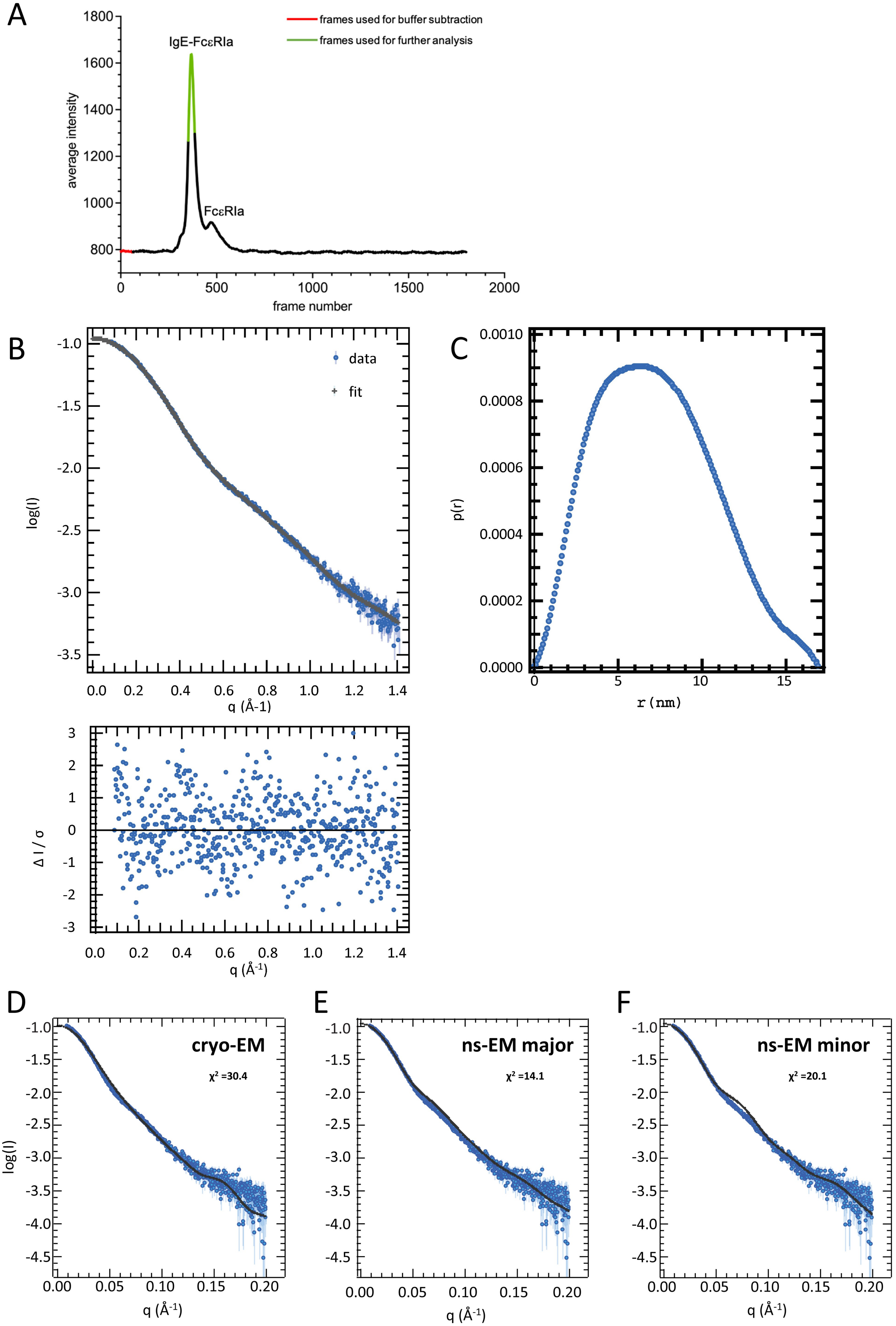
A
B
C
D
E
F
IgE-FcεRIa
FcεRIa
frames used for buffer subtraction
frames used for further analysis
average intensity
frame number
data
fit
log(I)
q (Å-1)
p(r)
r (nm)
Δ I / σ
q (Å-1)
cryo-EM
χ2 =30.4
ns-EM major
χ2 =14.1
ns-EM minor
χ2 =20.1

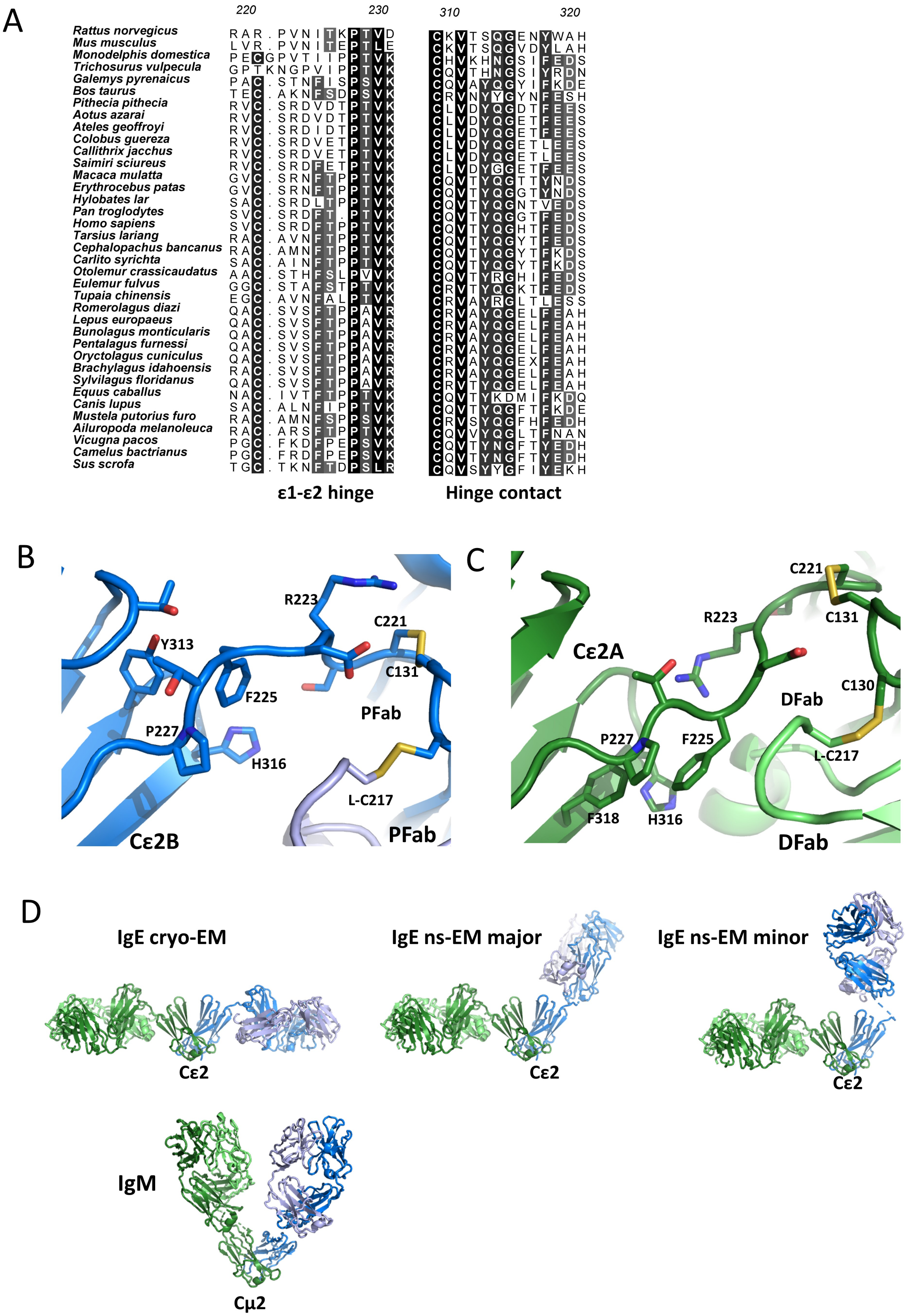
A
220
230
310
320
Rattus norvegicus
Mus musculus
Monodelphis domestica
Trichosurus vulpecula
Galemys pyrenaicus
Bos taurus
Pithecia pithecia
Aotus azarai
Ateles geoffroyi
Colobus guereza
Callithrix jacchus
Saimiri sciureus
Macaca mulatta
Erythrocebus patas
Hylobates lar
Pan troglodytes
Homo sapiens
Tarsius lariang
Cephalopachus bancanus
Carlito syrichta
Otolemur crassicaudatus
Eulemur fulvus
Tupaia chinensis
Romerolagus diazi
Lepus europaeus
Bunolagus monticularis
Pentalagus furnessi
Oryctolagus cuniculus
Brachylagus idahoensis
Sylvilagus floridanus
Equus caballus
Canis lupus
Mustela putorius furo
Ailuropoda melanoleuca
Vicugna pacos
Camelus bactrianus
Sus scrofa
ε1-ε2 hinge
Hinge contact
B
R223
C221
Y313
C131
F225
PFab
P227
H316
L-C217
Cε2B
PFab
C
C221
R223
C131
Cε2A
C130
DFab
P227
F225
L-C217
F318
H316
DFab
D
IgE cryo-EM
IgE ns-EM major
IgE ns-EM minor
Cε2
Cε2
Cε2
IgM
Cμ2

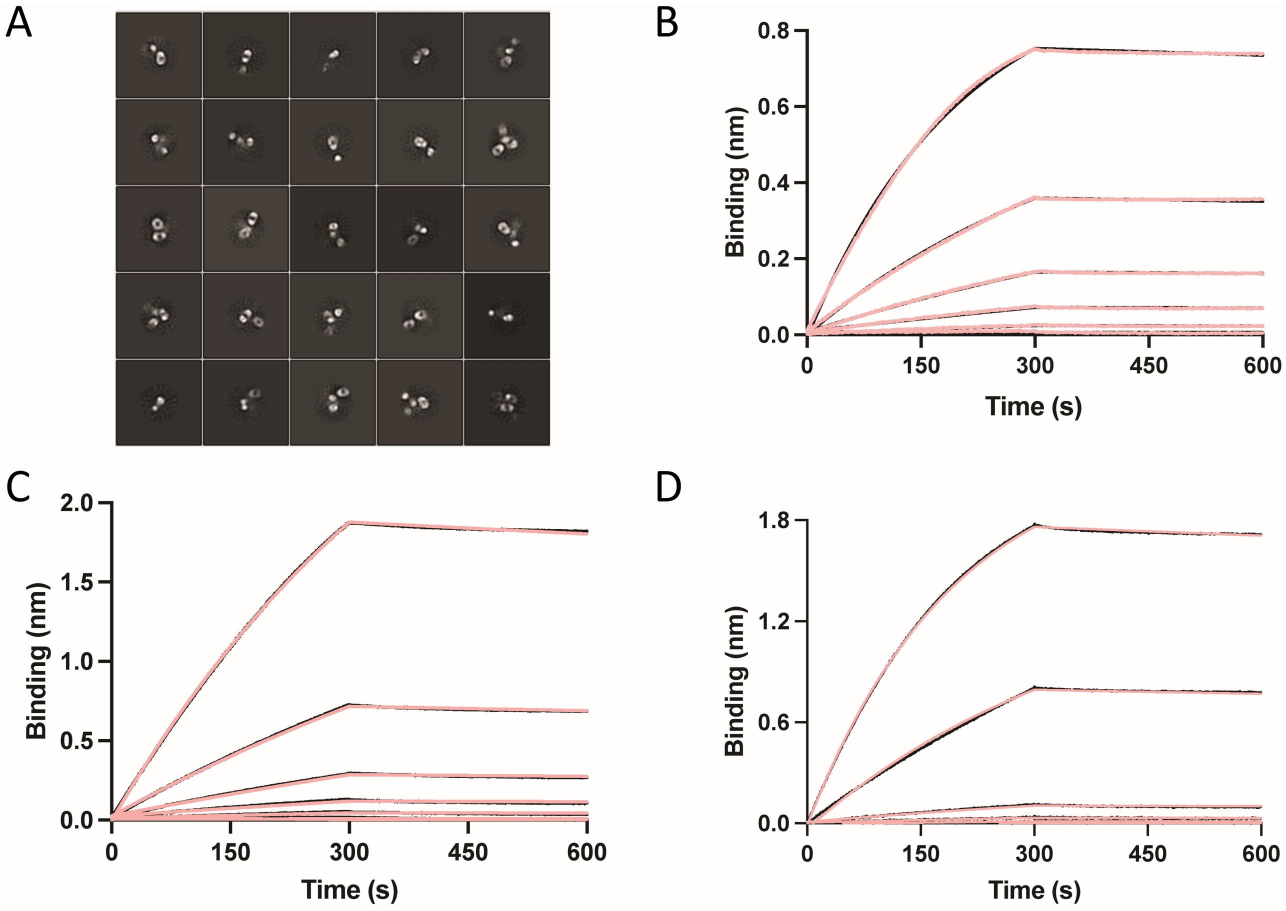


E

| | $k_{on}$ ($10^5$ $M^{-1}s^{-1}$) | $k_{off}$ ($10^{-5}$ $s^{-1}$) | $K_D$ (nM) |
|---|---|---|---|
| IgE Fc | 2.6 ± 0.43 | 8.2 ± 1.1 | 0.32 ± 0.06 |
| HMM5 | 1.2 ± 0.54 | 17 ± 6.4 | 1.4 ± 0.8 |
| HMM5L | 0.99 ± 0.12 | 11 ± 1.2 | 1.2 ± 0.2 |
